# Disease related changes in ATAC-Seq of more than 450 iPSC-derived motor neuron lines from ALS patients and controls

**DOI:** 10.1101/2023.09.11.557005

**Authors:** Stanislav Tsitkov, Kelsey Valentine, Velina Kozareva, Aneesh Donde, Aaron Frank, Susan Lei, the Answer ALS Consortium, Jennifer Van Eyk, Steve Finkbeiner, Jeffrey Rothstein, Leslie Thompson, Dhruv Sareen, Clive N. Svendsen, Ernest Fraenkel

## Abstract

Amyotrophic Lateral Sclerosis (ALS), like many other neurodegenerative diseases, is highly heritable, but with only a small fraction of cases explained by monogenic disease alleles. To better understand sporadic ALS, we report epigenomic profiles, as measured by ATAC-seq, of motor neuron cultures derived from a diverse group of 380 ALS patients and 80 healthy controls. We find that chromatin accessibility is heavily influenced by sex, the iPSC cell type of origin, ancestry, and the inherent variance arising from sequencing. Once these covariates are corrected for, we are able to identify robust ALS-specific signals in the data. Additionally, we find that the ATAC-seq data is able to predict ALS disease progression rates with similar accuracy to methods based on biomarkers and clinical status. These results suggest that iPSC-derived motor neurons recapitulate important disease-relevant epigenomic changes.

## INTRODUCTION

Amyotrophic lateral sclerosis (ALS)^1^ is a neurodegenerative disorder characterized by motor neuron loss. Its heritability has been estimated to be as high as 50%,^2^ but the known genetic factors account for less than 15% of cases. One possible explanation for the missing genetic component is that many diverse genetic causes lead to similar disruptions in pathways that are then exacerbated by non-genetic factors. Disease models based on induced pluripotent stem cell (iPSC) derived motor neurons generated from a broad cross-section of ALS patients may help identify such convergent, early effects. In this study, we examine the epigenomic profiles of more than five hundred cell cultures of iPSC-derived motor neurons (iMNs) generated from ALS patients and healthy controls to test for the presence of genetically driven, disease-relevant changes in chromatin accessibility and dysregulated transcriptional programs.

Epigenetics is an especially relevant level at which to look for genetically encoded ALS-specific impact in these cells. Changes in chromatin accessibility are generally attributed to the binding of pioneer transcription factors,^3^ DNA methylation, chromatin remodeling complexes, and histone post translational modifications (PTMs).^4^ Previous research has implicated several of these mechanisms in ALS pathology. For example, post-mortem spinal cord tissue from ALS patients exhibited elevated levels of the DNA methyltransferases DNMT1 and DNMT3A compared to controls.^5^ Motor neurons expressing FUS and TDP43 mutants exhibited a loss of subunits of the neuronal Brg1/Brm Associated Factor chromatin remodeling complex.^6^ Changes in the expression of the ALS genes *FUS*, *TDP43*, and *C9orf72* were found to be associated with changes in histone PTMs.^7^ Histone deacetylase inhibitors have even been proposed as a potential therapeutic for ALS.^8^ The identification of other ALS-specific epigenetic signatures will improve our understanding of early disease mechanisms and may suggest new therapeutic strategies. Because iPSCs undergo epigenetic reprogramming,^9^ the environmental contributions to ALS are likely to have been erased. As such, iPSC-derived cells allow a direct examination of the impact of as yet uncharacterized genetic factors on the epigenome.

The main problem in the identification of epigenetic signatures associated with ALS pathology is the heterogeneity in the genetic and clinical manifestations of the disease,^10^ and the scarcity of ALS patient-derived neuronal tissue. To address these problems, the Answer ALS consortium (AALS) is generating iPSC lines from the peripheral blood mononuclear cells (PBMCs) of over 800 ALS patients and 200 healthy controls that have been whole genome sequenced.^11^ The iPSC lines generated by AALS are differentiated into motor neurons (iMNs) and subjected to epigenomic, transcriptomic, and proteomic analysis. The advantage of iPSCs is that they can be generated from patient blood samples, grown in large quantities, and differentiated into disease-affected cell types.^12^ iPSC models of ALS have previously been used to characterize phenotypic patterns of neurodegeneration in mid-size cohorts of sporadic ALS patient iPSC-derived motor neurons derived from a population of Japanese ALS and control subjects,^13^ and to construct disease-associated protein-protein interaction networks for ALS cases associated with the mutant C9orf72 hexanucleotide repeat expansion.^14^ iPSCs are being used to model many other neurodegenerative diseases in smaller-scale studies and through large initiatives such as FOUNDIN-PD,^15^ iNDI,^12^ among others^12,16–25^.

Important technical challenges arise in studies of this scale, which necessarily have many sources of variation. The standard approach for analyzing omics data uses differential analysis with case status (ALS or healthy control) as the primary covariate. Such an approach is inappropriate in this setting. For example, sex imbalances in case/control groups can lead to false positive differential signals associated with the sex chromosomes. While sex is a covariate that can, in theory, be controlled for by meticulous study design or adjusted for in analysis, other covariates cannot be handled in these ways. Differentiated cell type composition, for example, was found to be the main driver of variation in AALS iMN gene expression, and it is not known until after the data are analyzed.^26^ The identification of robust ALS-associated signals requires a thorough understanding of sources of variation associated with sequencing, differentiation and clinical parameters.

In this study, we identify covariates that drive variation in the epigenomic profiles of 533 iPSC-derived motor neurons from ALS patients and healthy controls as measured by the Assay for Transposase-Accessible Chromatin using sequencing (ATAC-seq).^27^ This is one of the largest bulk ATAC-seq datasets generated by a single consortium (by bases sequenced), and the largest bulk ATAC-seq dataset overall from cell cultures of different donors using a single differentiation-protocol. The size of this dataset combined with the consistency of the data generation protocols allows it to be used as a tool to both investigate ALS-specific epigenomic signals, and establish practices for analyzing other ATAC-seq datasets. As might be expected from a study of a diverse population, conducted over several years, parameters such as sex, cell type composition, and sequencing efficiency drive much of the overall variance. Initially, we do not find any changes in chromatin accessibility when comparing all familial and sporadic ALS cases against controls – a finding that is also consistent with the variable etiology and phenotypes of ALS and in line with our previous study analyzing RNA-seq data in the identical patient groups. However, once these factors are accounted for, a strong differential signal is seen when stratifying by patients carrying the C9orf72 mutant hexanucleotide repeat expansion, a major risk factor for familial ALS. Surprisingly, we also find that the ATAC-seq data can be used to predict ALS progression rates at levels similar to clinical and neurofilament data. These results demonstrate that the chromatin accessibility of iPSC-derived motor neurons can reflect both genetic and clinical variation in ALS.

## RESULTS

### ATAC-seq data were generated for 533 motor neuron lines

ATAC-seq was conducted on 533 differentiated motor neuron lines from 460 unique donors (380 ALS patients and 80 healthy controls); 73 samples correspond to study design controls. The production of the motor neuron lines is described in detail in Baxi et al.^11^, and illustrated in brief in Figure 1a. Blood samples were collected from ALS patients and healthy controls. PBMCs, classified as either T-cells or non-T-cells (monocytes), were isolated from the blood samples and reprogrammed into iPSCs. The iPSCs were differentiated into motor neurons, and the resulting cell cultures were frozen and distributed across sequencing facilities. Variation between differentiation and sequencing batches was controlled for by including batch differentiation controls (BDCs) and batch technical controls (BTCs), and staining differentiated cell cultures with immunocytochemical (ICC) staining markers (Figure 1b, SI Section 1).

**Figure 1.**
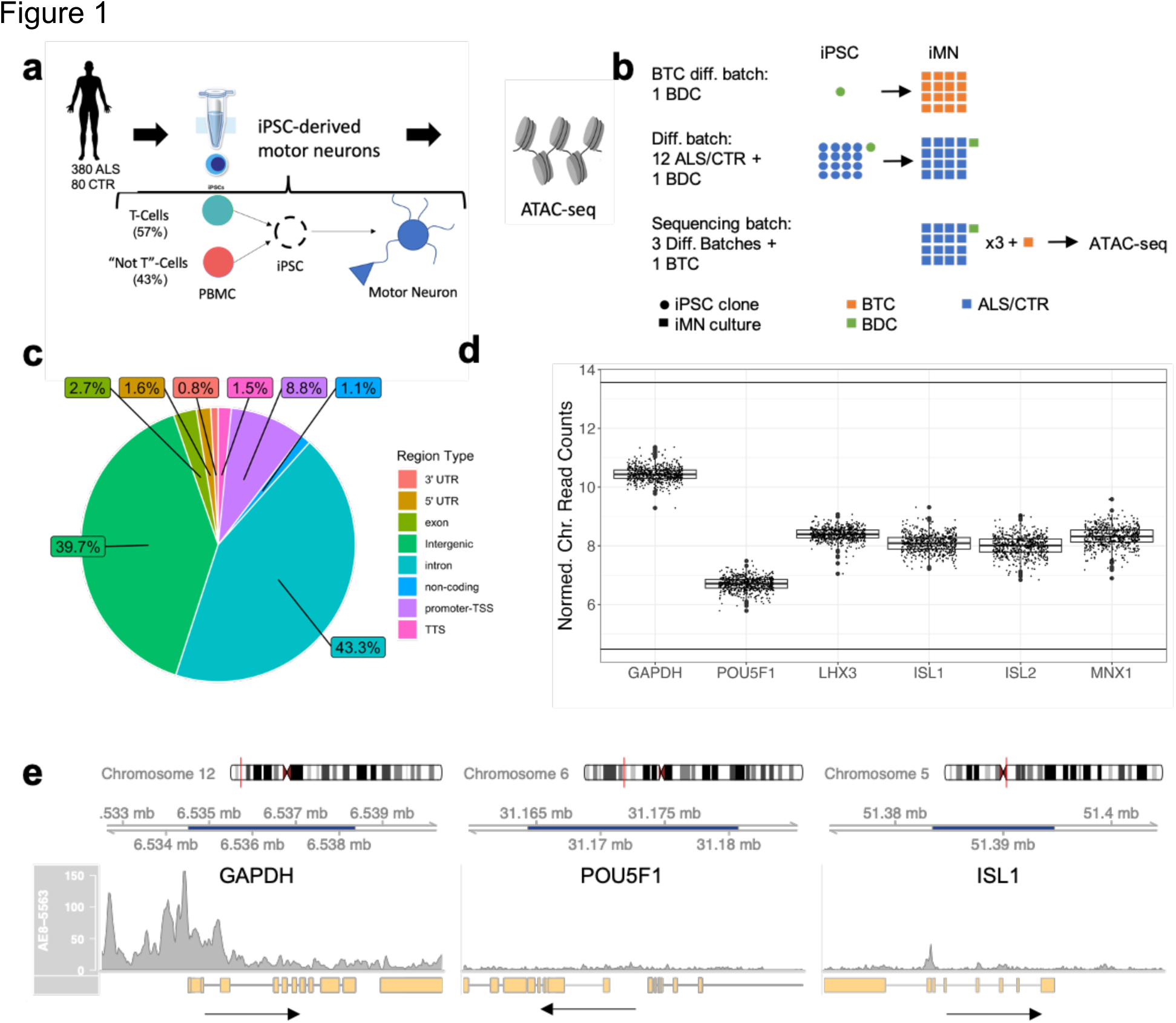
Answer ALS ATAC-seq data. **a**) Overview of AALS data generation protocol. PBMCs from ALS patients and healthy controls are reprogrammed into iPSCs, which are in turn differentiated into motor neurons and sent for sequencing. **b)** Overview of study design controls. Samples are divided into differentiation batches and sequencing batches. Each sequencing batch usually consists of three differentiation batches. A BDC is redifferentiated with each differentiation batch, and a BTC is resequenced with each sequencing batch. c) Pie chart showing distribution of region annotations. d) Normalized chromatin reads plotted for the promoter/TSS for the housekeeping gene, *GAPDH*, the pluripotency marker, *POU5F1*, and the spinal motor neuron-specific genes *LHX3*, *ISL1*, *ISL2*, and *MNX1*. e) Raw read coverage plots spanning the gene bodies of *GAPDH*, *POU5F1*, and *ISL1*. Dark blue shading of the genome axis scale indicates the gene body location. All plots are drawn on the same scale. Arrows point in the direction of transcription.

### Evaluation of ATAC-seq data quality

Overall, ATAC-seq-specific alignment QC metrics satisfied ENCODE guidelines (see Figures S1a-c, Methods).^28^ The consensus peak set contained 100,363 chromatin regions, of which approximately 80% were intronic/intergenic, and 10% were promoters/5’ UTRs (Figure 1c). To evaluate the quality of the ATAC-seq data as it pertains to motor neurons, we examined the chromatin accessibility of genes specific to motor neurons as done previously by Sahinyan et al.^29^. Chromatin accessibility was assessed for the housekeeping gene, *GAPDH*, a set of spinal motor neuron-specific genes (*LHX3*, *ISL1*, *ISL2*, *MNX1*),^30,31^ and as a negative control, the pluripotency marker *POU5F1*.^29,32^ As expected, out of the six genes tested, only the promoter/TSS chromatin region for *POU5F1* was not accessible (Figure 1d,e). Data reproducibility was confirmed by the assessment of inter-sample correlations and comparisons to re-differentiated samples (SI Section 2).

### Most variably-accessible regions are driven by three sources of variation

To investigate the underlying factors contributing to variance in the data, we conducted principal component analysis (PCA) (Figure 2a, Figure S2a) and found three major sources of variance: sex, iPSC cell type of origin, and sequencing instrument (SI Section 3). In fact, we found that applying UMAP to the top 100 MVARs separated samples into four distinct clusters defined by sex and PBMC type (Figure 2b). We used this UMAP representation to estimate the PBMC type for samples with missing PBMC labels in downstream analyses. Additionally, we noted that the BTC/BDC samples completely separated from the remainder of the female samples (Figure 2a) and found that the separation is partially driven by genomic variants specific to the BTC/BDC samples; for example, a genomic structural variant characterized by a 2 kb deletion dramatically affects chromatin accessibility in BTC/BDC samples (SI Section 4).

**Figure 2.**
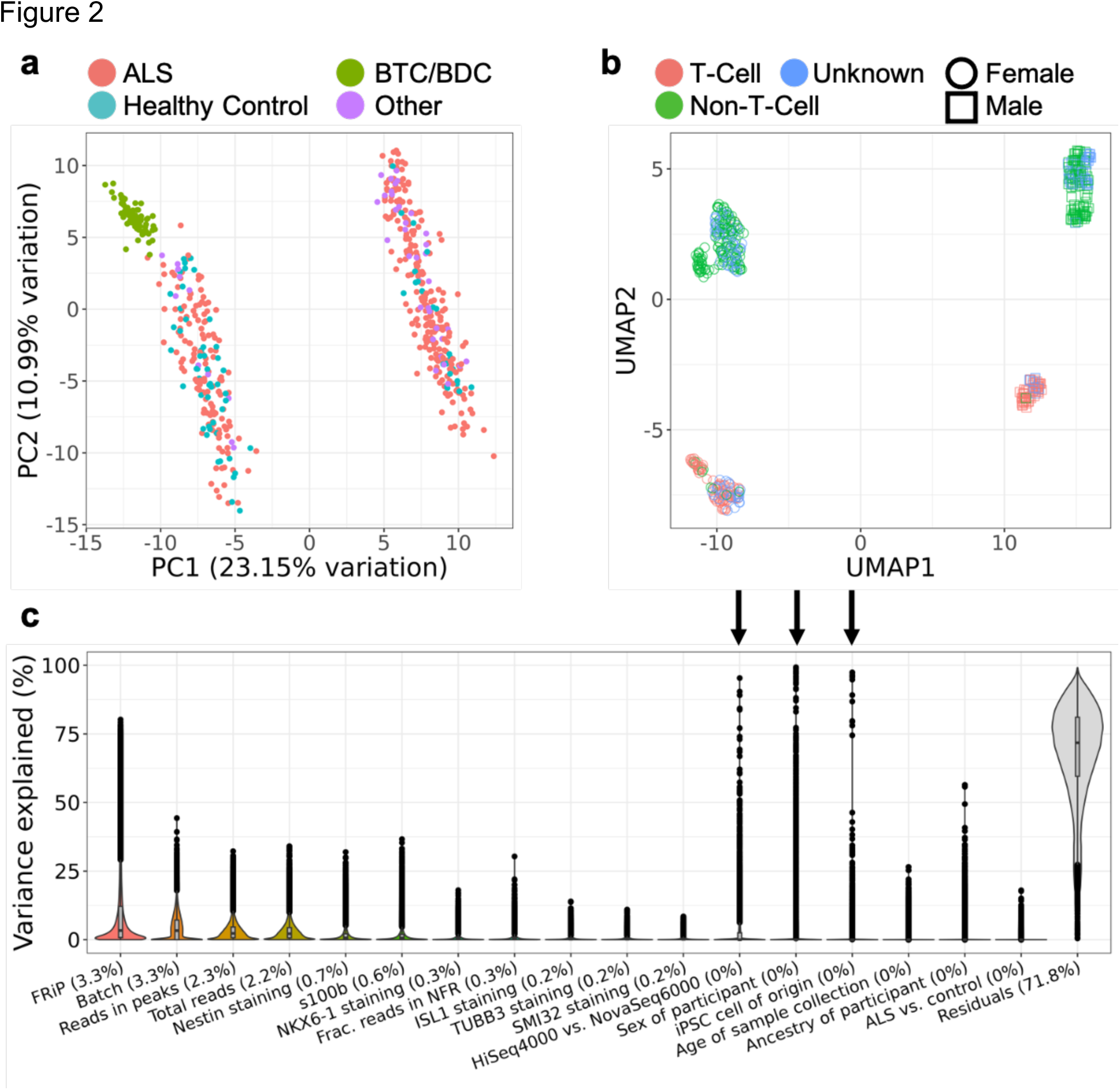
Drivers of variation in chromatin accessibility. **a**) Biplot of PC1 and PC2 from principal component analysis on top 500 MVARs including all samples. “Other” refers to samples from individuals with non-ALS motor neuron disease and asymptomatic ALS. **b)** UMAP applied to 100 MVARs separates samples into clusters by sex and PBMC type. **c)** Explained variance in chromatin accessibility by selected covariates across all chromatin regions. Data was generated by fitting a linear mixed effects model to normalized chromatin reads for each chromatin region. Percentages indicate median contribution to variance. Arrows indicate covariates that were found to drive variation in the PCA of the 500 MVARs.

To explore the effects of a more comprehensive set of covariates, we additionally fit the chromatin accessibility of each region to a set of 16 covariates using a linear mixed effects model (Figure 2c). These covariates reflected the sequencing, differentiation, clinical, and demographic aspects of each sample. Differentiation batch explained the most variance across all regions, but with few regions explaining over 25% of the variance, indicating a small effect size. The association of several regions with the sequencing-associated covariates, Fraction of Reads in Peaks (FRiP) score and sequencer, could be explained by the normalization methods used and raw read length (SI Section 5). The ICC staining markers contributed to the variation of the samples in a manner similar to that found in the gene expression data, with the most variation driven by S100B and Nestin; however, these markers had a small effect size overall, explaining less than 25% of the variance for all but 32 and 16 regions, respectively. As expected from the PCA analysis, the variability of several regions was driven by sex, PBMC type, and sequencer. Interestingly, we also found a dependence on ancestry, which was not observed in gene expression.^26^

### Differentiation-associated covariates

It was interesting that unsupervised clustering separated samples along PBMC type (T-cell or non-T-cell; Figure 2b); in the RNA-seq data, this was only observed when the gene set was constrained to four T-cell receptor associated genes.^26^ Examining DARs associated with PBMC type, we found that regions located next to the T-cell receptor delta anti-sense 1 (*TRD-AS1*) gene were dramatically less accessible in T-cell derived cell lines as would be expected (Figures 3a-b) due to T-cell receptor rearrangements. Beyond the *TRD-AS1* regions, there are 180 DARs associated with PBMC type (FDR < 0.01, abs(log2FC) > 0.5); 25 of these regions are annotated as promoters. The top 5 significant promoter regions, other than those that that correspond to pseudogenes or lincRNAs, are labeled in Figure 3a. To confirm that the differential signal was not a sequencing artifact, we compared the accessibility of these regions to the gene expression for matched samples, and found high correlations (Figures 3c-f). The existence of genes not associated with the T-Cell receptor loci whose promoter accessibility and expression are dependent on PBMC type illustrates a modest, but detectable, impact of epigenetic memory in our dataset: the chromatin accessibility and gene expression profiles of differentiated cell cultures depend on the initial PBMC type. To determine whether the observed epigenetic memory could be attributed to differentiation bias resulting in different cell type distributions, we compared PBMC type to ICC staining data of the motor neuron cultures. There were no significant associations (p<0.01) between PBMC type and the percent of cells that stained positive for S100B, Nestin, ISL1, NKX6.1, TUJ1, or SMI32. In general, chromatin accessibility was less correlated with ICC staining markers than gene expression (SI Section 6). This indicated that the observed epigenetic memory, which influences a small set of genes, cannot be attributed to differentiation bias.

**Figure 3.**
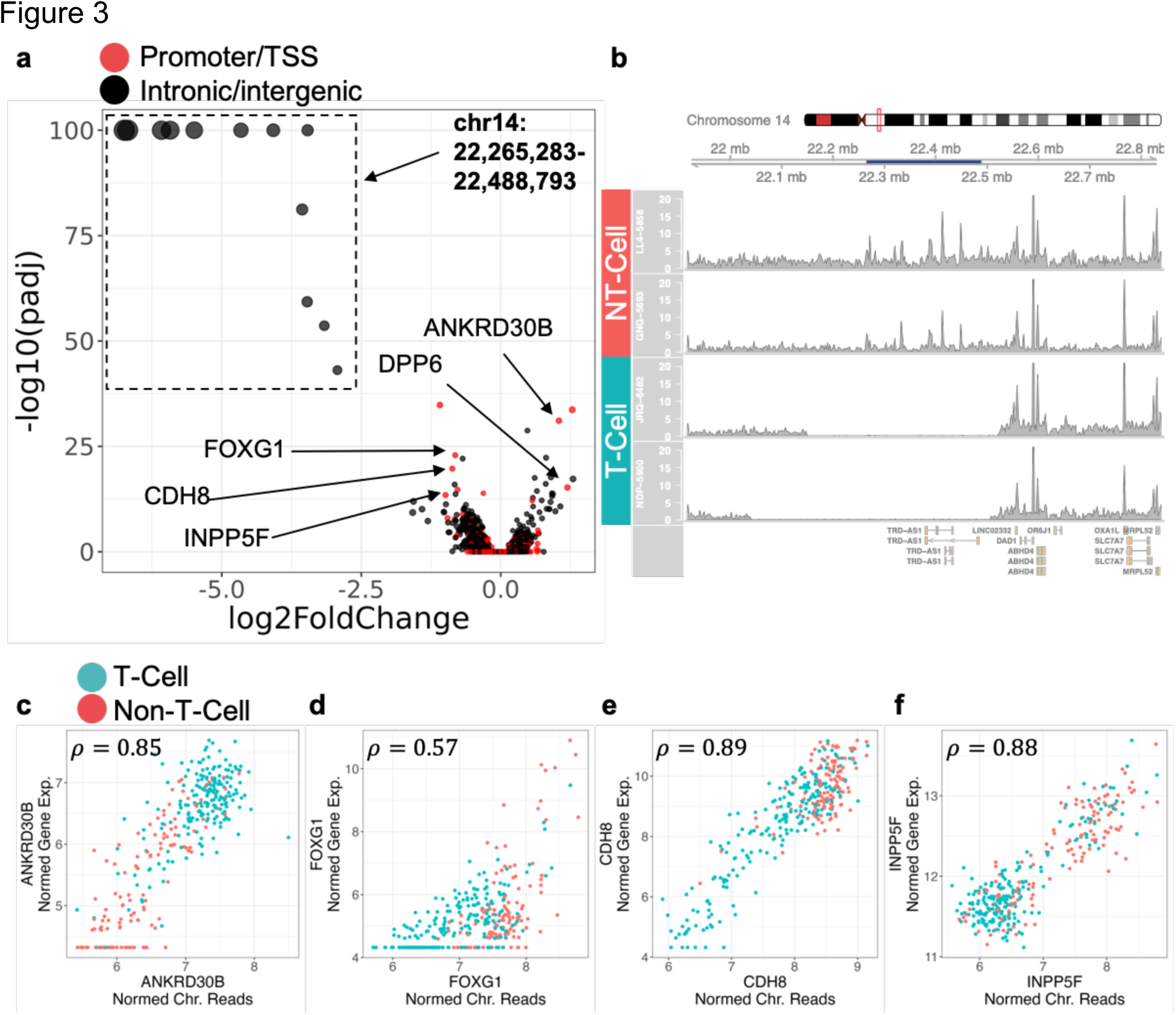
Differentiation-associated covariates. **a**) Volcano plot for PBMC-associated differential signal. **b)** Coverage plot spanning T-cell receptor genomic region for two T-cell-derived samples and two non-T-cell-derived samples. Dark blue region in genome axis scale spans the regions in the box from (a). **c-f)** Plots of chromatin accessibility against gene expression for selected genes from (a). Samples colored according to PBMC type. Pearson correlations between plotted chromatin accessibility and gene expression are indicated in top left corner.

### Clinical and demographic covariates

The key question in the analysis of iPSC derived motor neurons is whether omic profiles capture clinical information about the individuals from whom the cells were derived. In Figure 2c, we found that out of the four patient-specific covariates tested, only sex and ancestry drove variance in chromatin accessibility; the contributions of case status and age were negligible. The lack of an age-associated signal is not surprising, as iPSC-reprogrammed cells exhibit elongated telomeres, reduced oxidative stress, and a loss of senescence markers, all hallmarks of younger cells.^33–35^ The ancestry signal was confirmed with differential analysis and revealed 47 DARs (adj. p-val. <0.01, Figure S4a), of which 16 were more accessible in individuals of African ancestry. The top DAR by p-value was the promoter/TSS of *RNF135* (adj. p-value 3e-14, log2FC = 0.8) (Figure 4a); despite the presence of six mismatches to the reference genome in the aligned reads (Figure S4b), subjects of African ancestry were found to have a higher accessibility of this promoter region than subjects of European ancestry. These mismatches are consistent with the genomics data (rsIDs: rs7221217, rs7221238, rs7219775, rs7221473, rs7225888, rs7211440).^11^ Interestingly, the promoter/TSS of *RNF135* has previously been identified to exhibit hypomethylation in subjects of African ancestry, which is consistent with our observation of lower accessibility.^36^ The remaining two clinical covariates, sex and ALS status, are examined in the following sections.

**Figure 4.**
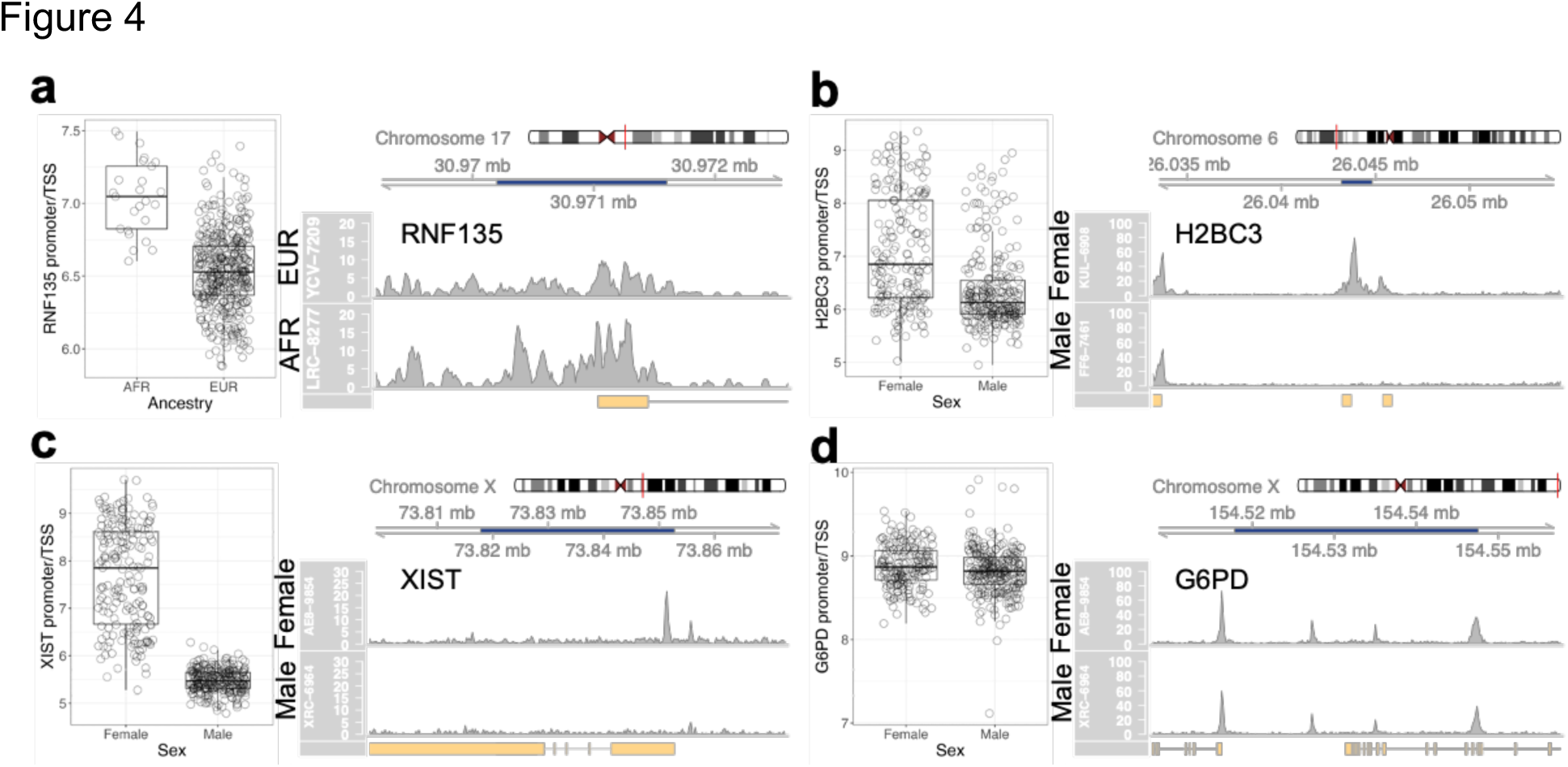
Clinical covariates. **a**) (Left) Normalized read counts and (right) example coverage plot for individuals of African (AFR) and European (EUR) ancestry for the promoter/TSS of *RNF135* (adj. p-value = 3-14, log2FC = –0.78). **b)** (Left) Normalized read counts and (right) example coverage plot for *H2BC3*, a sex-associated autosomal DAR (adj. p-value = 3e-17, log2FC = –1.16). **c)** (Left) Normalized read counts and (right) example coverage plot for *XIST*, a sex-associated DAR that escapes X-inactivation (adj. p-value < 1e-278, log2FC = –4.00). **d)** (Left) Normalized read counts and (right) example coverage plot for *G6PD*, a housekeeping gene on chromosome X (adj. p-value = 0.02, log2FC = –0.05).

### Sex-associated DARs are not limited to sex chromosomes and reveal X-chromosome inactivation

There were 72 significant DARs associated with sex (adj. p-value <0.01, abs(log2FC)>1); of these regions, 40 were Y-chromosomal, 22 were X-chromosomal, and 10 were autosomal (Figure S4c). The only promoter in the 10 autosomal significant DARs was that of *H2BC3*, a histone encoded on chromosome 6 (Figure 4b). The most significant DAR on chromosome X corresponded to the promoter of XIST, a gene responsible for X chromosome inactivation (Figure 4c). The most significant DAR on chromosome Y corresponded to the promoter of *ZFY* (Figure S4d). All of these genes were also differentially expressed; for example, the *XIST* promoter accessibility had a correlation of 0.94 with *XIST* gene expression, and the accessibility of the *H2BC3* region was most highly correlated with the expression of its corresponding gene (0.89), the histone *H3C3* (0.72), and the histone *H4C9* (0.46). The remaining autosomal regions were all most highly correlated with the expression of genes on the Y chromosome. Motif enrichment of the 10 significant autosomal DARs using HOMER^37^ did not return any significantly enriched motifs. The inactivation of the X chromosome was confirmed by examining numbers of background reads (SI Section 7).

### C9orf72 TSS is differentially accessible in ALS patients with C9 hexanucleotide repeat expansion

After accounting for all previously mentioned covariates, there were no significant DARs associated with ALS case status (Figure 2c). We hypothesized that the heterogeneity of ALS might obscure disease-relevant signals. To test this hypothesis, we compared ALS cases that were known to harbor the mutant *C9orf72* hexanucleotide repeat expansion (n=31: C9+) to verified C9-negative cases and healthy controls (n=116: HC). For both the C9+/C9– and C9+/HC comparisons, the *C9orf72* TSS had significantly lower normalized chromatin reads (p.adj = 1e-50) in C9+ ALS patients, with a consistent log2FC of –0.6 (30% decrease in accessibility) (normalized counts and coverage plot in Figure 5a). The differential signal extends over 6 raw read lengths away from the repeat expansion, indicating that the signal is not a mapping artifact (Figure S5a). Interestingly, the inclusion of the FRiP score as a covariate in the differential analysis improved the adjusted p-value from 1e-20 to 1e-50, but did not influence the log2FC. The observation of lower chromatin reads is consistent with the hypothesis of haploinsufficiency of the *C9orf72* transcript contributing to neurodegeneration and agrees with our previous report showing reduced *C9orf72* transcript levels.^26,38^ We did not observe any significant dependence between C9 repeat length and chromatin reads that could not be explained by C9 status.

**Figure 5.**
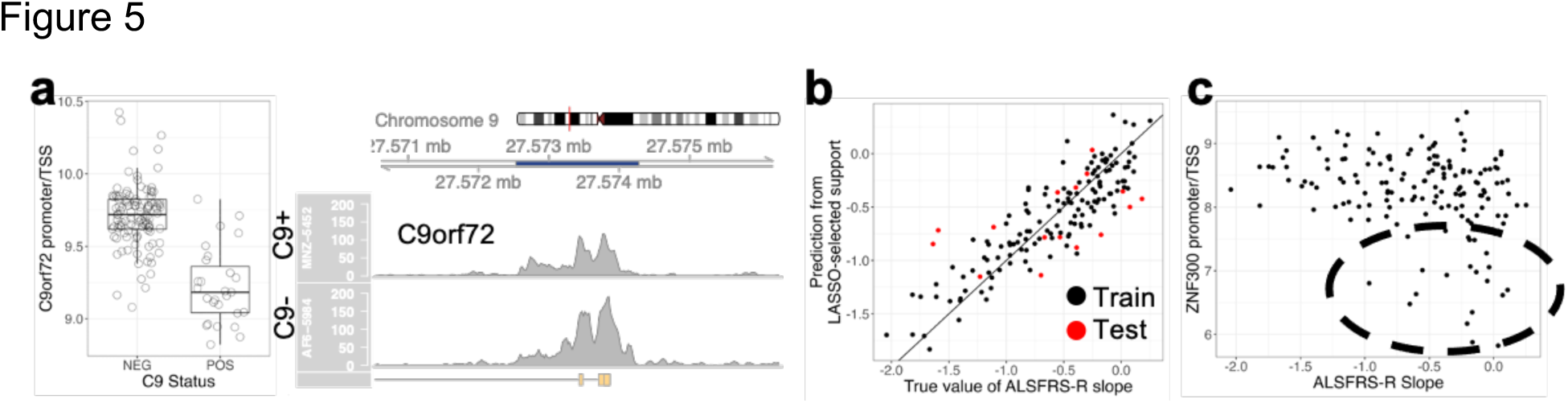
AALS ATAC-Seq ALS signals. **a**) (Left) Normalized chromatin read counts and (right) example coverage plot for the promoter/TSS of *C9orf72*; the coverage plot corresponds to an ALS case with a repeat expansion length of 274 (C9+) and an ALS case without the C9orf72 mutation (C9-). **b)** Prediction vs true value of ALSFRS-R slope; samples used to train the classifier are black, and samples used to test the classifier are red. Black line is a reference line of slope 1 and intercept 0. **c)** Normalized chromatin read counts for the promoter of *ZNF300*, a chromatin region associated with ALSFRS-R slope from the classifier; a set of samples that appear to both have slower disease progressions and low accessibility are circled.

### ATAC-Seq signals predict ALSFRS-R slope

Rates of disease progression in ALS are highly variable, with the time from first symptoms until death ranging from months to decades. One of the most widely used measures of the rate of disease progression is the linear slope of the ALSFRS-R score across time. To explore whether chromatin accessibility contains information related to progression, we sought to predict ALSFRS-R slopes from ATAC-Seq.

ALSFRS-R slope was not significantly associated with the accessibility of any chromatin region. This is not necessarily surprising, as any genetic component to the rate of progression is likely to be multifactorial in nature. To take this into account, we used linear regression with a LASSO penalty to search for a small set of regions that were the most predictive. LASSO linear regression with ten-fold cross-validation on the training set resulted in a model consisting of 27 regions (Table S1). On the training data (140 samples, see Methods), the model had a root mean square error (RMSE) of 0.23 and a correlation between predicted and true ALSFRS-R slope of 0.79. When the fit predictor was used to estimate the ALSFRS-R slopes of the testing data (16 samples, see Methods), it resulted in an RMSE of 0.46 and a correlation of 0.53 (Figure 5b). Surprisingly, performance on the held-out test data are on par with the models that have attempted to predict ALSFRS-R slope from clinical data; a model using neurofilament concentrations at diagnosis exhibited RMSEs of 0.4 and 0.9 in validation cohorts,^39^ and models based on clinical metadata returned root mean squared deviations of 0.54.^40^

The genes in the regions selected by the model (Table S1) include several of interest. *CHCHD2* is a gene definitively associated with Parkinson’s disease and closely related to the ALS-associated *CHCHD10*, and *LMX1A* is a transcription factor involved in neurogenesis and also associated with Parkinson’s disease. Another transcription factor in the feature set is *ZNF300*; in a plot of ALSFRS-R slope against the normalized chromatin accessibility of the *ZNF300* promoter/TSS, it appears that a subset of ALS cases with slower progression rates also exhibit lower *ZNF300* promoter/TSS chromatin accessibility (Figure 5c). Additionally, the chromatin accessibility of this gene has a correlation of 0.89 with its gene expression. Previous work has shown that *ZNF300* is a transcription repressor that localizes to the nucleus and is expressed in the heart, skeletal muscle, and brain.^41^ It is also associated with NF-kB pathway activation and MAPK/ERK signaling.^42^ The NF-kB pathway has been suggested to play a role in ALS disease progression, suggesting that the lower chromatin accessibility of the *ZNF300* TSS potentially indicates a protective mechanism against ALS progression.^43^

## DISCUSSION

With over 5 trillion bases sequenced, the ATAC-seq data presented in this work is the largest ATAC-seq dataset generated for iPSC motor neurons to date, and it is one of the largest ATAC-seq datasets generated by a single consortium overall. The consistency of the biological sample being produced (i.e., motor neuron cultures from hundreds of individuals rather than mixed tissues from the same individual) makes this dataset amenable to revealing insights that extend beyond ALS disease-associated signatures to other covariates, such as sex and iPSC tissue of origin.

### Challenges with producing consistent ATAC-seq data over multiple years

To assemble a dataset of this size, it is necessary to conduct a study which spans years. Over this period, the goal of maintaining consistent data generation methods can come into conflict with facility changes, instrumentation modernizations, and other unavoidable events. This can in turn influence downstream processing results. We showed an example of this in the analysis of the sequencer-associated differential signal. In general, the best way to monitor these changes is to examine batch-specific QC metrics. In future studies, efforts should be focused not only on surpassing a specific set of QC metrics, such as those defined by ENCODE, but also to ensure that the final quality control metrics have minimal batch to batch variance.

### Genetic variants and mismapping

Chromatin read counts are susceptible to influence by genetic variants and the resulting mismapping. The best example of this is the apparent difference in chromatin accessibility between the line used as a BTC/BDC and other control lines. We determined that the difference is due to a 2kb deletion, which we confirmed through comparisons with the genomics data. This signal arises despite a read mapping rate above 97.5% across almost all samples. We benefited from the fact that full-genome sequences were available for each sample in our study. In the future, we recommend that differential accessibility signals are verified against genomic data of the same sample in inter-individual comparisons, or that raw reads are aligned to individualized genomes when those data are available. The latter approach has previously been shown to alter peak calls in ChIP-seq data and alignment in RNA-seq data.^44^ When genomics data is not available, inspection of raw reads aligned to differential peaks could reveal SNPs; genome coverage visualization software such as Gviz^45^ provides a streamlined approach for this.

### ATAC seq normalization

Several studies have examined the question of which normalization approach is superior for the analysis of ATAC-seq data. Our analysis was based solely in the framework provided by DESeq2, and we decided to use the default DESeq2 geometric median of ratios algorithm to estimate normalization factors. We found that it performed similarly to normalizing by reads in peaks (RiP). Notably, it outperformed normalization by total reads, which failed to identify the *C9orf72* TSS DAR in C9+/C9-ALS comparisons and to generate separation by PBMC type in PCA. However, RiP normalization alone is imperfect; indeed, the covariates total reads, reads in peaks, and FRiP score are closely related, and we observed a strong dependence of certain chromatin regions on FriP score even after RiP normalization. These strong dependences are a concern as they will lead to false positive results if co-accessibility analyses, such as WGCNA, are used. Future work could focus on the analysis of alternative methods for normalization, which could include data from other omics modalities as validation.

### Epigenetic memory in iPSC-derived motor neuron cultures

The concept of epigenetic memory describes the phenomenon wherein iPSC-derived cells retain epigenetic characteristics of the cell type from which the iPSC clone was dedifferentiated.^46^ In the analysis of this data, we observed that the chromatin accessibility of several regions was significantly associated with the PBMC type. A fraction of the affected regions corresponded to T-cell receptor loci, where genomic TCR rearrangements prevented reads from mapping. These DARs are therefore not reflective of epigenetic memory.^47^ At the same time, the differential signal at several other chromatin regions could not be explained by mapping artifacts. For example, the TSS for *FOXG1* is significantly more accessible in non-T-cell derived samples than it is in T-cell derived samples, while the TSS for *ANKRD30B* exhibits the opposite dependence on PBMC type. *FOXG1* is a neurodevelopmental factor that functions as a transcriptional repressor, promotes neurogenesis, and inhibits gliogenesis; mutations in the gene are associated with Rett’s syndrome.^48^ *ANKRD30B* is a gene that is expressed in the brain; it has recently been found to be differentially methylated in subsets of patients with Alzheimer’s disease^49^ and Williams Syndrome.^50^

### Different populations of cell types

The ATAC-seq data exhibited significant correlations with ICC staining markers, but these signals were not as strong as those found in the RNA-seq data, and they mainly corresponded to intronic/intergenic chromatin regions. This was surprising because the epigenome is responsible for establishing cellular phenotypes.^51^ There are a few possible explanations for the weaker signal compared to the RNA-seq data. First, there is more biological noise in the transduction of an epigenomic signal into a proteomic signal, as it requires both transcription and translation. Another possible explanation for the discrepancy is that the expression of ICC markers is a response to an external stimulus, which could induce changes in the cellular populations of transcription factors. For example, S100B can be released by damaged cells.^52^ Finally, it is conceivable that the same ICC staining markers can stain multiple cell types, all of which have unique epigenomic signatures, resulting in low correlations with chromatin accessibility.

### Demographic differences

We identified both ancestry– and sex-associated differential signals in the ATAC-seq data. The observation of ancestry-specific differential accessibility in iPSC-derived motor neurons highlights the need to explore whether ancestry may play a role in motor neuron function and survival.

The sex associated DARs spanned both the autosomal and sex chromosomes, a finding which is consistent with previous work. For example, sex has been found to influence autosomal chromatin accessibility in immune cells in an age-dependent manner.^53^ We showed that the differential signal associated with chromosome X was mostly driven by background reads from the inactivated chromosome X, with the exception of 22 DARs that clearly escaped inactivation and including the promoter/TSS for XIST. It was important to establish X chromosome dosage compensation in these iPSC lines, as its erosion has been found to be a limitation in iPSC-based disease modeling.^55,56^ While low passage iPSCs retain X inactivation, longer culture times lead to gradual re-activation of the inactivated X chromosome that is not reversed by differentiation.^57^ Hallmarks of eroded dosage compensation include decreased *XIST* gene expression and a loss of H3K27me3 marks, and can lead to the remodeling of the iPSC proteome.^57,58^ It is also interesting to note that in gene expression data, the number of sex-associated differentially expressed genes (78 genes, abs(log2FC)>1, adj. p-value<0.01) was much higher than in the ATAC-seq data; similar to the observations with ICC staining markers, this suggests that there is an additional level of regulation governing the expression of these genes that is not apparent at the epigenome level. Overall, the fact that there is still sex-based variant gene expression and chromatin accessibility at the level of the motor neuron cultures raises questions regarding whether sex may impact motor neuron function and survival.

### ALS subtypes should guide search for disease signatures

There were no global differences between ALS cases and controls, and the strongest ALS-specific signal revealed by the ATAC-seq data was a difference in chromatin accessibility at the *C9orf72* TSS for C9+ ALS cases. This finding is perhaps not surprising. There is strong evidence that ALS is not one disease, but several different diseases that culminate in the same clinical phenotype of upper and lower motor neuron degeneration.^1^ There are more than 25 known independent genetic causes of ALS, which collectively explain less than 15% of cases. Thus, it is likely that the variability among ALS patients may be greater than any common “ALS signature.” In addition, due to epigenetic reprogramming, iPSCs are likely to best represent early phases of disease. Thus, these samples may reflect the diverse early causes of the disease and not later stages of cell death that may be common to more patients. The iPSC data in this study, therefore, are best used to explore how genomic factors beyond the known disease-causing loci contribute to the high heritability of ALS.

### ALSFRS-R slope

Several studies have attempted to use clinical data and other biomarkers to group ALS patients and predict ALS disease progression. The Prize4Life challenge crowdsourced machine learning models to predict ALS disease progression from clinical data, identifying time from disease onset, ALSFRS, forced vital capacity, and blood pressure among the top predictors.^40^ Semi-supervised machine learning models applied to clinical data of Italian ALS patients was found to separate ALS patients according to the Chio criteria.^59^ The distribution of T cell populations in the CSF of ALS patients was found to be associated with ALS disease progression.^60^ In this study, we show that the ATAC-seq data of iPSC-derived motor neurons from ALS patients has a predictive power for disease progression rate that is on-par or better than predictions from clinical data or blood-based biomarkers.

We seek to answer a different question than these prior studies. We asked whether iPSC-based models retain clinically relevant signals. On the one hand, the high heritability of ALS suggests that they should. On the other hand, iPSC-derived motor neurons are expected to exhibit the characteristics of a ‘younger’ cell, with reversed senescence due to reprogramming.^33^ Are iPSC-derived neurons too ‘young’? Our results suggest that ALS-relevant genomic influences emerge very early in this system. It remains an open question how early such signals might emerge in patients, but some studies of presymptomatic C9ORF72 carriers suggest that some effects may be detectable early in life.^61,62^

Much more work will be needed to identify how the genetic variants influence disease progression. By their nature, the machine learning models we used cannot not determine which correlated features are causal, only which have the strongest predictive value in a particular dataset. Nevertheless, it is interesting that the chromatin regions returned by the ALSFRS-R slope predictor are associated with neurodegenerative diseases. For example, *LMX1A* and *CHCHD2* have been previously associated with Parkinson’s disease. We also identify the potentially protective role of decreased accessibility at the *ZNF300* promoter/TSS. Finally, it is likely that ALSFRS-R may not be the best signal to try and predict, as it is a sum of scores reflecting deficits in extremely diverse symptoms and may mask important variability among patients with the same overall score. Future work will need to use more sophisticated analyses of clinical states and will need to integrate other omic signals.

## CONCLUSIONS

In this study, we examined the epigenomic profiles of one of the largest sets of iPSC derived motor neurons generated to date. Whereas these cell lines were generated to identify ALS-specific disease signatures, we showed that the epigenomic analysis of the iMN cultures could be used to gain insights that extend beyond the disease. We found that chromatin accessibility measurements were influenced by clinical covariates, such as sex and ancestry, differentiation-associated covariates, such as the iPSC cell type of origin, and sequencing-associated covariates, such as FRiP score and the read length used. Importantly, as this data is used by a wider audience, these covariates must be factored into any differential and co-accessibility analyses to avoid false-positive signals that are associated with ALS case status.

We described two ALS signals in this study. The first one was a decrease in the chromatin accessibility of the *C9orf72* promoter/TSS in samples exhibiting the mutant hexanucleotide repeat expansion. This supports the hypothesis of haploinsufficiency for the *C9orf72* transcript contributing to disease.^38^ Additionally, we found that the epigenomics data could be used to construct a predictor of ALSFRS-R slope, and identified the downregulation of the *ZNF300* gene as having a potentially protective effect.

Overall, this paper underscores the value of conducting large-scale investigations of iPSC-derived cells for the study of ALS. After carefully compensating for sources of variance, these data reveal some of the complex interplay between chromatin accessibility, genetics, and disease subtypes, including, surprisingly, an association between epigenomic signals and the rate of disease progression. The expanding multi-omic data from Answer ALS and other efforts raises the prospect that integration of these data with other omics modalities and ALS omics datasets will uncover new directions in ALS research and help identify novel therapies.

## METHODS

### Generation of iPSC motor neuron cultures

Methods for generating iPSCs, quality control, and iPSC generation were described previously in Baxi et al.^11^ and Workman et al.^26^

### ATAC-Seq experimental methods and quality control

As we wrote in Baxi et al.,^11^ ATAC-seq sample prep, sequencing and peak generation were carried out by Diagenode Inc. as further described.87 Briefly, cells were lysed in ATAC-seq resuspension buffer (RSB; 10 mM Tris-HCl, pH 7.4, 10 mM NaCl, 3 mM MgCl2 and protease inhibitors) with a mixture of detergents (0.1% Tween-20, 0.1% NP-40 and 0.01% digitonin) on ice for 5 min. The lysis reaction was washed out with additional ATAC–RSB containing 0.1% Tween-20 and inverted to mix. Then 50,000 nuclei were collected and centrifuged at 450 r.c.f. for 5 min at 4 °C. The pellet was resuspended in 50 μl of transposition mixture (25 μl of 2× Illumina Tagment DNA buffer, 2.5 μl of Illumina Tagment DNA enzyme, 16.5 μl of phosphate-buffered saline, 0.5 μl of 1% digitonin, 0.5 μl of 10% Tween-20 and 5 μl of water). The transposition reaction was incubated at 37 °C for 30 min followed by DNA purification. An initial PCR amplification was performed on the tagmented DNA using Nextera indexing primers (Illumina). Real-time (RT)-qPCR was run with a fraction of the tagmented DNA to determine the number of additional PCR cycles needed, and a final PCR amplification was performed. Size selection was done using AMPure XP beads (Beckman Coulter) to remove small, unwanted fragments (<100 bp). The final libraries were sequenced using the Illumina NextSeq platform (PE, 75-nt kit). All samples passed QC checks that included morphological evaluation of nuclei, fluorescence-based electrophoresis of libraries to assess size distribution and RT-qPCR to assess the enrichment of open chromatin sites.

### ATAC-seq read alignment and peak calling

ATAC-Seq data was processed using the ENCODE-DCC ATAC-seq pipeline v1.7.1. Reads were aligned to GRCh38 genome build using Bowtie2. The quality of the sequencing was assessed using FastQC. Samples had 62.0 +/− 0.8 (s.e.) million total reads after mitochondrial filtering and deduplication, with 90% of samples having over 40 million reads (Figure S1a). The average sample FRiP score was 0.242 +/− 0.002 (s.e.), with 77% of samples having a FRiP score higher than 0.2 (Figure S1b). The average transcription start site enrichment (TSSE) was 14.2 +/− 0.1 (s.e.), with 99% of samples having a TSSE greater than 7 (Figure S1c). All samples had a distinct nucleosome free-region in fragment length distribution plots. We identified open chromatin regions separately for each sample using the peak-calling software MACS2 and determined differentially open sites using DESeq2 (FDR < 0.1).

### Generation of consensus peakset and raw counts matrix

After peak calling, using the R package DiffBind,^63^ a consensus peakset was constructed by retaining peaks that were open in at least 10% of samples; it consisted of 100,363 variable–width chromatin regions. These regions were annotated using HOMER^37^. Chromatin read counts from the consensus peakset were normalized using the DESeq2^64^ *vst* function. A parametric fit was used for the dispersion estimate, and the default DESeq2 geometric median of ratios was used for the scaling factor estimate.

### Evaluation of processed data quality

Coverage plots and read pileups were generated using the R package *Gviz*^45^ (Figure 1e). Sample-wise Pearson correlations within the BTC, BDC, and inter-individual groups (Figure S1d) were calculated using the columns of the normalized counts matrix. Outlying samples were identified using complete link hierarchical clustering with Euclidean distance on the columns of the correlation matrix (heatmap for BDCs shown in Figure S1f).

### Replication cohort analysis

23 samples were redifferentiated, sequenced and compared to the initial cohort. To compare samples in the replication cohort to the initial cohort, Diffbind^63^ was run on the 46 total samples from the initial and replication cohorts to generate a new consensus peakset and raw counts matrix. The raw counts matrix was normalized using the DESeq2 *vst* function as before. After normalization, an inter-sample correlation matrix was constructed by calculation Pearson correlations between individual samples. Samples were clustered by applying complete link hierarchical clustering on the Euclidean distance between the columns of this correlation matrix.

### PCA/UMAP analysis

PCA was conducted on the top 500 MVARs using the R package PCAtools.^65^ The UMAP representation on the 100 MVARs was generated using the R package UMAP.^66^ For downstream analysis, the UMAP representation was used to estimate the PBMC type identity of samples with missing data. PCA in Figure 2a was conducted while including BDC/BTC samples, and PCA in Figures S2c-g excluded BDC/BTC samples.

### Genomics data analysis

Genomics vcf files were obtained from Baxi et al.^11^ and analyzed using Pysam.^67^

### Fitting linear mixed model

Each row (chromatin region) of the normalized read counts matrix was fit to a set of 16 covariates with a linear mixed model using the R package variancePartition.^68^ Discrete variables (differentiation batch, sequencer, sex, case status, PBMC type, ancestry) were modeled with random effects and continuous variables were modeled with fixed effects.

### Differential analysis for generating volcano plots

Differentially accessible regions (DARs) associated with covariates were identified by running DESeq2 on the raw counts matrix using a parametric dispersion fit and the default size parameter estimates while including FRiP score, sequencer, sex, PBMC type, and ALS case status as covariates. The p-values were adjusted for multiple hypothesis testing using the Bonferroni correction. Adjusted p-values and log2FC values for each covariate were used for volcano plots.

### Simulations to identify the role of sequencer in influencing differential accessibility results

The effect of raw read length on differential accessibility was evaluated on a subset of 20 samples (10 with 75bp HiSeq4000 reads, and 10 with 50bp NovaSeq6000 reads). Raw reads were trimmed down to 50bp and 36 bp lengths, and realigned to the reference genome. To generate a consensus peakset and raw counts matrix, Diffbind was run on a set of 50 samples: 10 75bp HiSeq4000 samples, 10 trimmed 50 bp HiSeq4000 samples, 10 trimmed bp HiSeq4000 samples, 10 with 50bp NovaSeq6000, and 10 trimmed 36bp NovaSeq6000. The same set of samples were used for each trimming. The raw counts were analyzed using DESeq2 vst normalization and differential analysis.

The differential signal in this test set recapitulated the signal from analyzing all samples, with 75 bp HiSeq4000 samples exhibiting higher measured accessibility across most DARs (Figure S3a). Following this confirmation, the 75bp reads were trimmed down to 50bp, and the analysis was repeated; the most significant DARs had lost their significance (Figure S3b). We found that the measured chromatin accessibility of the most significant DARs fell with decreasing read length (Figure S3c).

### Correlations with gene expression

There were 335 samples that had both gene expression and chromatin accessibility reported. Generation of raw counts for gene expression as described in Workman et al.^26^ Raw counts from gene expression data were normalized using the DESeq2 *vst* function and the default DESeq2 parameters (parametric fit for the dispersion estimate, geometric median of ratios for size factor estimate).

### Estimating reads not in peaks in X chromosome

The total number of reads that mapped to the X chromosome was estimated from the mitochondrial filtered, deduplicated BAM files using Samtools.^69^ The raw reads in peaks from the X chromosome was estimated using the Diffbind-output raw counts matrix. The number of reads not in peaks (RniP) on the X chromosome was estimated by subtracting these two quantities. The X chromosome RniP was normalized by dividing by the total number of reads to generate Figure S4e.

### LASSO linear regression

The predictor was constructed on a set of filtered ALS cases. In total, there were 242 ALS samples with a recorded ALSFRS-R slope. The normalized chromatin read counts matrix was first subset to chromatin regions within 2 kb of a TSS and samples with a recorded ALSFRS-R slope. An inter-sample correlation matrix was constructed using this subset counts matrix.

Euclidean distance complete-link hierarchical clustering on this correlation matrix revealed a set of 12 samples that were poorly correlated with other samples; these samples were excluded from further analysis (Figure S5b). In the remaining sample set, six samples were identified as having an outlying ALSFRS-R slope and excluded as well (Figure S5c). Outliers were defined in the traditional way as samples that exhibited an ALSFRS-R slope that was more than 1.5 interquartile ranges away from the first and third quartiles. To mitigate the effects of batch-to-batch variation, samples that came from batches with less than 4 representatives in the sample set were removed; this caused the removal of 68 samples. Next, samples were randomly divided into a 90/10 split for a training set (140 samples) and testing set (16 samples). LASSO linear regression was conducted using the R package glmnet.^70^

