## Supplementary Information for "Disease related changes in ATAC-Seq of more than 450 iPSC-derived motor neuron lines from ALS patients and controls"

[1]Department of Biological Engineering, Massachusetts Institute of Technology, Cambridge, MA, USA. [2]Cedars-Sinai Biomanufacturing Center, Cedars-Sinai Medical Center, Los Angeles, CA, USA. [3]Advanced Clinical Biosystems Research Institute, Cedars-Sinai Medical Center, Los Angeles, CA, USA. [4]Center for Systems and Therapeutics, Gladstone Institutes, University of California, San Francisco, San Francisco, CA, USA. [5]Taube/Koret Center for Neurodegenerative Disease, Gladstone Institutes, University of California, San Francisco, San Francisco, CA, USA. [6]Departments of Neurology and Physiology, University of California, San Francisco, San Francisco, CA, USA. [7]Brain Science Institute, Johns Hopkins University School of Medicine, Baltimore, MD, USA. [8]Department of Neurology, Johns Hopkins University School of Medicine, Baltimore, MD, USA. [9]Institute for Memory Impairments and Neurological Disorders, University of California, Irvine, CA, USA. [10]Department of Biological Chemistry, University of California, Irvine, CA, USA. [11]Department of Neurobiology and Behavior, University of California, Irvine, CA, USA. [12]Department of Psychiatry and Human Behavior and Sue and Bill Gross Stem Cell Center, University of California, Irvine, CA, USA. [13]The Board of Governors Regenerative Medicine Institute, Cedars-Sinai Medical Center, Los Angeles, CA, USA.

[+]Corresponding Author

Contact:

Ernest Fraenkel


**Table of Contents**

Supplementary Information Section 1: **Batch control design** 3

Supplementary Information Section 2: **Reproducibility analysis** 3

Supplementary Information Section 3: **PCA analysis** 4

Supplementary Information Section 4: **Example of genomic variant in B(D/T)C sample** 4

Supplementary Information Section 5: **Sequencing-associated covariates** 5

Supplementary Information Section 6: **Correlations between chromatin accessibility and the S100B staining marker** 7

Supplementary Information Section 7: **Confirmation of X-chromosome inactivation** 8

Supplementary Information Figure S1: **Alignment QC and replicates** 9

Supplementary Information Figure S2: **Drivers of variation in most variably accessible regions** 10

Supplementary Information Figure S3: **Influence of sequencer on chromatin accessibility** 11

Supplementary Information Figure S4: **Clinical and demographic covariates extended data** 13

Supplementary Information Figure S5: **AALS ATAC-seq ALS signals** 14

**References** 15

Supplementary Information Section 1

### Batch control design

The scale of the study required that patient samples be batched for both differentiation and sequencing. A maximum of 12 lines were differentiated in each batch. These were collected into larger batches of 22-36 lines for sequencing. Accordingly, several quality control (QC) measures were incorporated into the study design. Variation in differentiated cell type composition was evaluated by staining cell cultures with six immunocytochemical staining (ICC) markers: S100B, ISL1, NKX6-1, TUJ1, SMI32, and Nestin. Variation between differentiation batches was controlled for by re-differentiating the same iPSC line with each differentiation batch (the batch differentiation control, or BDC). Variation between sequencing batches was controlled for by resequencing the same sample with each sequencing batch (the batch technical control, or BTC) (Figure 1b). Both BDCs and BTCs were differentiated from the same female healthy control iPSC clone.

Supplementary Information Section 2

### Reproducibility analysis

While replicate guidelines have not been defined for bulk ATAC-seq, almost all samples fell within the ENCODE guidelines for bulk RNA-seq replicates. Most isogenic replicates (BDCs) had a Pearson correlation greater than 0.9, and most anisogenic replicates (inter-individual) had a Pearson correlation greater than 0.8 (Figure S1d). The removal of a limited number of outlying samples (3 out of 55 BDCs, 2 out of 18 BTCs, and 5 out of 493 inter-individual) improved these metrics considerably (Figure S1e, f). As a final quality control measure, 22 iPSC clones were re-differentiated, sequenced, and compared to samples in the initial cohort. Notably, 21 out of 22 replicates from the same iPSC clones clustered together when correlating samples by the 500 most variably accessible autosomal chromatin regions (MVARs) (Figure S1g). Bringing the sample correlation results together, we concluded that the data was highly reproducible.

Supplementary Information Section 3

### PCA analysis

In a PCA of the 500 MVARs, sex is the primary source of variance, separating samples along PC1 (Figure S2b); this separation is driven, as expected, by regions on the Y chromosome.  If the sex chromosomes are excluded from the analysis, samples separate by PBMC type (Figure S2c), i.e. whether the iPSC clone of the corresponding sample was derived from a T-cell or a non-T-cell (monocyte, non-T-cell).  If samples are further stratified by PBMC type, there is separation along PC1/2 that corresponds to the sequencing instrument used (HiSeq4000 or NovaSeq6000) (Figs. S2d-e).

Supplementary Information Section 4:

### Example of genomic variant in B(D/T)C sample

In a PCA of the top 500 MVARs across all samples, we noted that the BTC/BDC samples separated from the remainder of the female samples (Figure 2a).  This separation is partially driven by genomic variants specific to the BTC/BDC samples.  For example, BTC/BDC samples differ from healthy controls in an intergenic differentially accessible chromatin region (DAR) on chromosome 17 (adjusted p-value <1e-100) (Figure S2f).  Differential accessibility for this DAR was confirmed by examining raw read coverage (Figure S2g).  Genomic analysis revealed that the BTC/BDC iPSC clone has a rare genomic structural variant, which is characterized by a 2kb deletion at this DAR, that was previously catalogued by Abel et al. in their characterization of 17,795 human genomes (locations of catalogued deletions are shown in red track in Figure S2g).[1]

Supplementary Information Section 5

### Sequencing associated covariates

Several regions were highly associated with the Fraction of Reads in Peaks (FRiP) score, an ATAC-seq QC metric that reflects the signal to noise ratio of ATAC-seq data; correlations between region accessibility and FRiP score were found to be as high as 0.9 (METTL8 promoter/TSS), and as low as -0.7 (KMT2C intron). This suggested that the chosen normalization strategy using reads in peaks (RiP) did not completely remove sequencing depth associated signals (see Discussion) and led us to include FRiP score as a covariate in downstream differential analyses.

*Change in sequencer from HiSeq4000 to NovaSeq6000 affected chromatin accessibility*
As the result of the modernization of the sequencing facility, the instrument used for sequencing the ATAC-seq libraries changed from the HiSeq4000 to the NovaSeq6000 after the first 20 of 50 total differentiation batches (160 samples). This also resulted in a shortening of the raw sequencing read length from 75 bp to 50 bp. Comparing samples from the different platforms revealed 131 regions that differed significantly (adjusted p-value < 0.1, abs(log2FC)>1) (Figure S3a), with 118 being more accessible on the HiSeq4000 instrument (example in Figure S3b). However, there were also regions that were much more accessible when sequenced on the NovaSeq6000 (example in Figure S3c). We carried out simulations to determine the effect of read length on differential accessibility (see Methods, Figures S3d-e). Measured accessibility of the most significant DARs fell with decreasing read length (Figure S3d-f); this effect is consistent with the expectation for a low sequence complexity region. Surprisingly, one chromatin region that seemed to have significantly *lower* measured accessibility for the 75 bp HiSeq4000 samples became more accessible when reads were trimmed (Figure S3g). The raw reads that mapped to this region originally mapped to the mitochondrial genome when 75 bp long, but were mis-aligned to (non-mitochondrial) chromosome 1 when 50 bp long. A pileup of the raw reads that map to this region reveals a large number of consistent base-pair mismatches which are not observed in the genomics data for this region (Figure S3h). Based on these results, we conclude that the most significant sequencer-associated DARs are raw read length-dependent mapping artifacts. Nevertheless, while trimming reads reduced the log2FC of the most differentially accessible regions, it did not have a significant impact on overall differential analysis results. There was a similar number of DARs with a low effect size (abs(log2FC)<1) (Figs. S3d-e); both the p-value and log2FC remained similar for most genes

(Figs. S3i-j) and were largely accounted for by our corrections for FRiP and TSSE (Figure S3k-l). Since read trimming was only confirmed to eliminate a small number of DARs, we used the untrimmed data in our subsequent analyses.

Supplementary Information Section 6:

**Correlations between chromatin accessibility and the S100B staining marker**

The associations between chromatin accessibility and ICC staining markers were notably weaker than those in gene expression data. Nevertheless, we did find that the percent of cells that stained positive for S100B (percent S100B) significantly correlated with the chromatin accessibility of several regions. Percent S100B was most highly correlated with the accessibility of an intron of *VPS13D* ($\rho$=0.67). The promoter region most correlated to percent S100B corresponded to that of *WBP1L* ($\rho$=0.66). Interestingly, the gene expression of *WBP1L* has a correlation of only 0.50 with percent S100B. A similar effect was observed in the RNA-seq data, where the expression of the *S100B* gene was less correlated with percent S100B than several other genes. This suggests that there is an additional level of regulation beyond gene expression and chromatin accessibility that drives the percent S100B-associated signal in this ATAC-seq data.

Supplementary Information Section 7

### Confirmation of X-chromosome inactivation

A source of variability in the use of iPSC-derived cell models of disease is the erosion of X chromosome inactivation which can lead to imbalanced gene expression.[2] This is especially important if the disease has an X-linked component, and ALS has been characterized to be more prevalent in males.[3] So, our goal was to determine whether the X chromosome was inactivated. On the one hand, the chromatin region corresponding to the housekeeping gene, *G6PD*, which is on the X chromosome, exhibits the same accessibility between males and females (Figure 4d). Beyond this, the volcano plot (Figure S4c) identified a distinct set of 22 regions that likely escape inactivation, exhibiting a log2FC greater than 1; they corresponded to the promoter/TSS of *XIST* (Figure 4c), and the introns of *DANT1/2* and *FIRRE*. The gene *XIST* is a known escape gene. On the other hand, there are over 1000 regions on the X chromosome that exhibit a significantly higher measured accessibility in females but with a abs(log2FC)<1. For these regions, we hypothesized that slightly elevated signal in females was due to background reads from the inactivated X chromosome. Indeed, we found that background reads on the X chromosome were 50% higher in female samples than male samples, as measured by reads *not* in peaks normalized to total reads across all chromosomes (Figure S4e). Because this higher baseline would also result in inflated reads in peaks, we concluded that the X chromosome had indeed been inactivated, consistent with previous observations of gene expression.[4]

Supplementary Figure S1

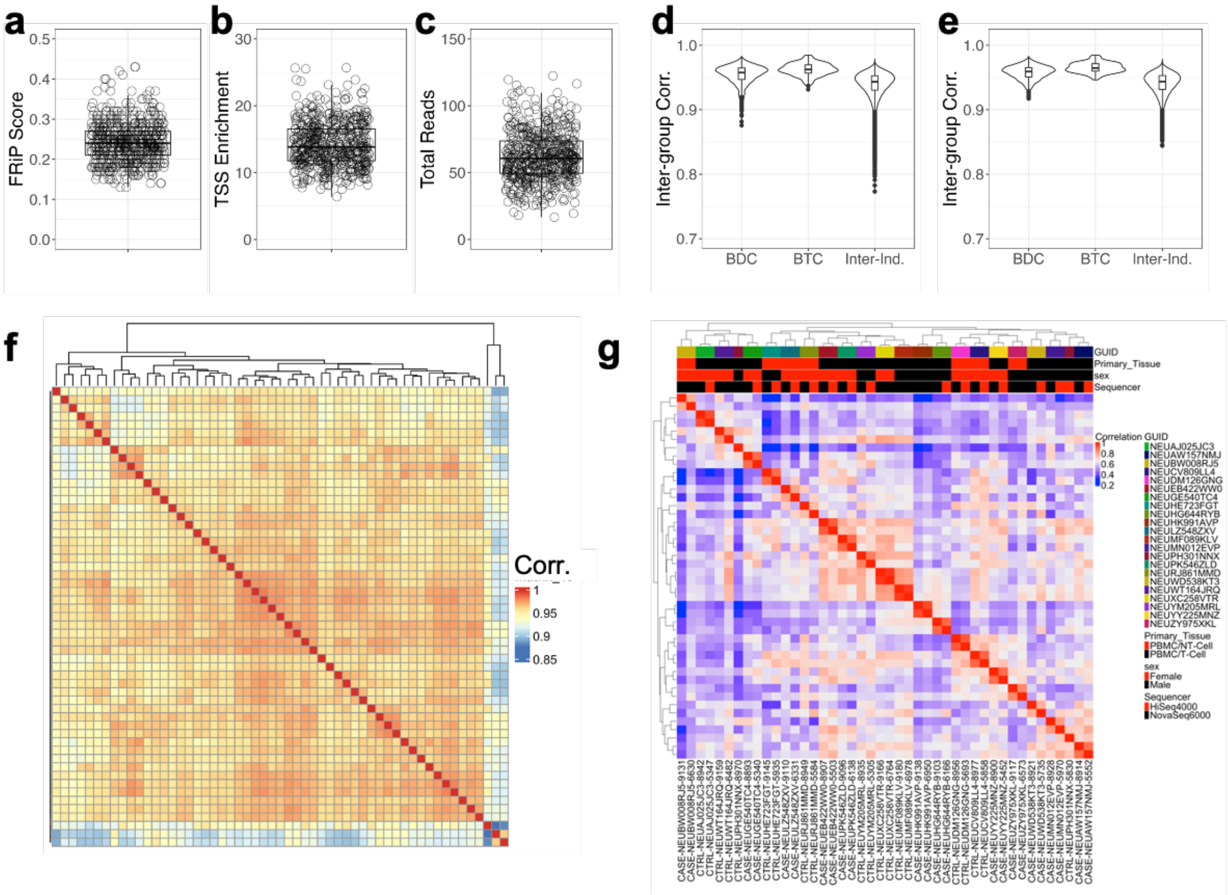


**Supplementary Figure S1. Alignment QC and replicates. a-c)** Distributions of alignment QC metrics across all samples. Total reads refers to total reads (in millions) after mitochondrial filtering and deduplication. d**)** Distributions of Pearson correlation coefficients between BDC samples, between BTC samples, and between all non-B(T/D)C samples. e**)** Same plot as (d) but with outlying samples, as identified by hierarchical clustering in (f) removed. **f)** Heatmap of correlation coefficients between BDC samples shows three outliers (last three columns). **g)** Heatmap of correlation coefficients between differentiation replicates for 500 MVARs; 21 out of 22 samples cluster together when correlating across top 500 MVARs. Regions on the X and Y chromosomes are not included in the analysis.

Supplementary Figure S2

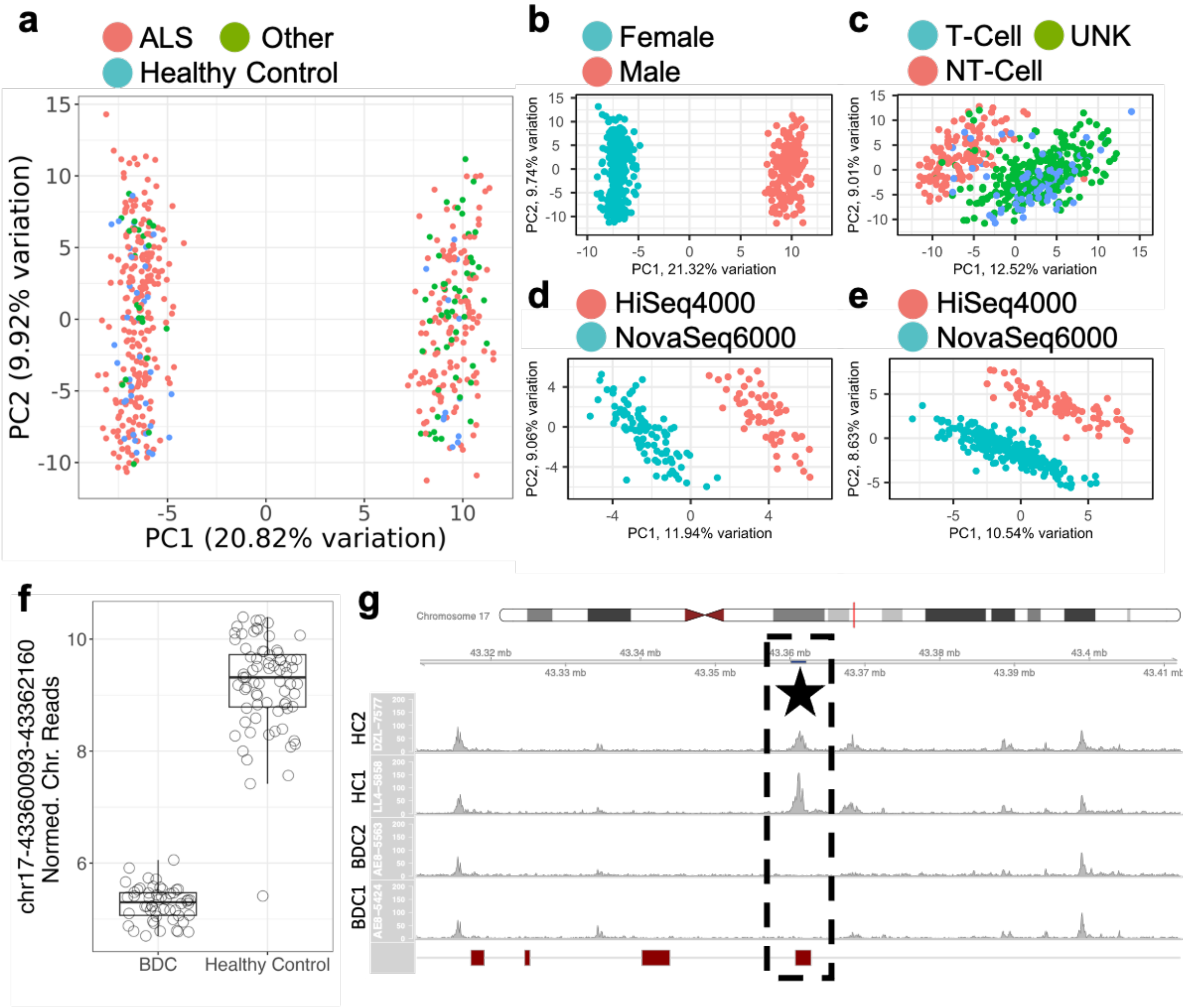


**Supplementary Figure S2. Drivers of variation in most variably accessible regions. a)** Biplot of PC1 and PC2 when PCA is conducted on 500 MVARs after removing B(T/D)C samples. **b)** Same as (a) but with points labeled by sex. **c)** PC1/PC2 biplot of PCA conducted on 500 MVARs after removing sex chromosomes. **d-e)** PC1/PC2 biplot of PCA conducted on 500 MVARs after removing sex chromosomes and stratifying by PBMC type. Left, non-T-cell, right, T-cell. **f)** BDC samples exhibit a significantly differentially accessible region (DAR) at an intergenic position on chromosome 17 (adj. p-value < 1e-100). **g)** Raw read coverage plots of the DAR from (f) for 2 BDC samples and two healthy controls (HC). Genomic position of DAR is highlighted in dark blue in the genome axis scale (above the star) and surrounded by the dashed box. Red track at the bottom corresponds to structural genomic variants identified by Abel et al.[1]

Supplementary Figure S3

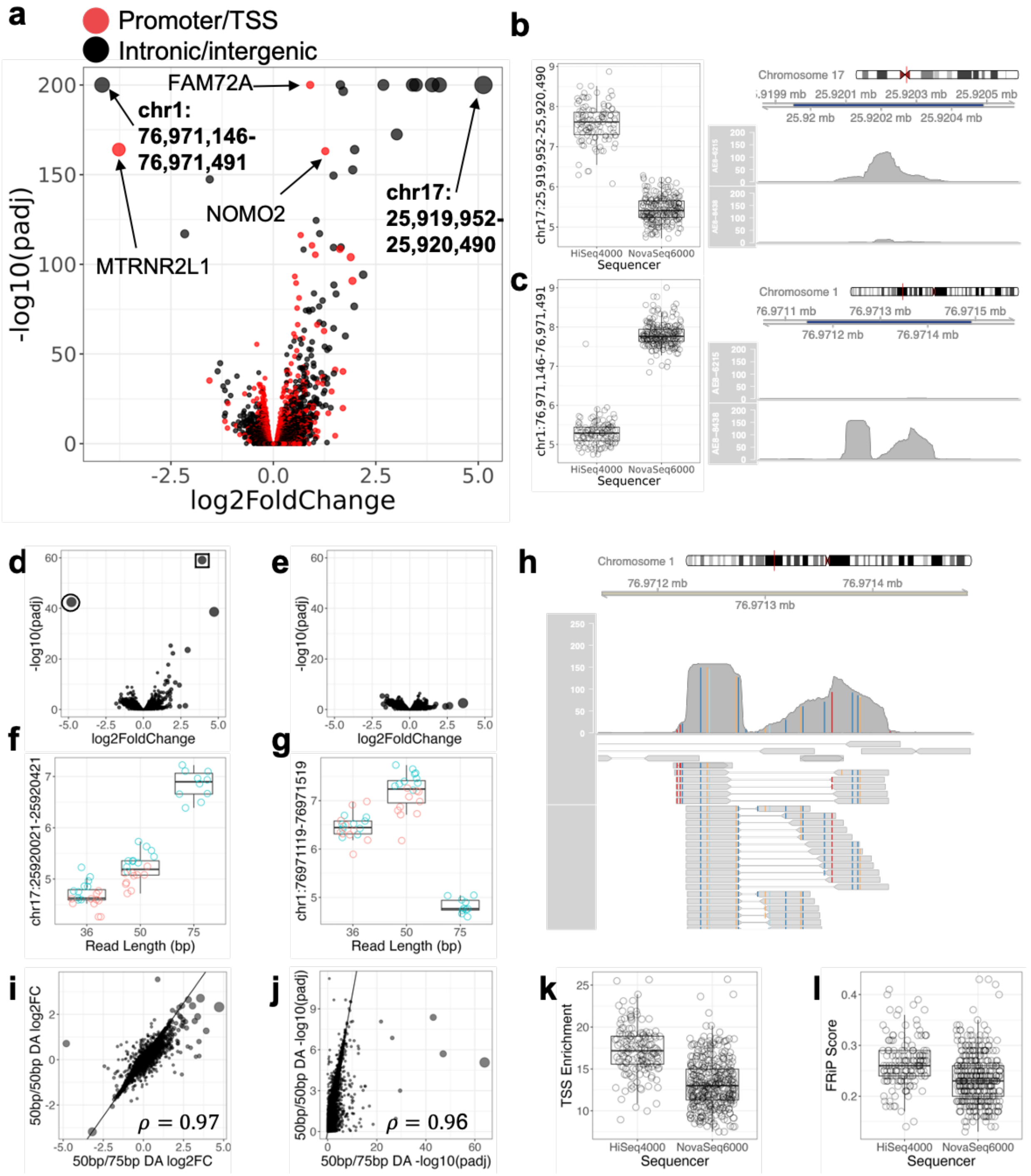


**Supplementary Figure S3. Influence of sequencer on chromatin accessibility. a)** Volcano plot of sequencer-associated differential signal. Point sizes reflect higher and more significant effect sizes. **b)** Normalized chromatin read counts (left) and coverage plot (right) for a BTC sample that was sequenced on the HiSeq4000 (top), and NovaSeq6000 (bottom) for the most differentially accessible region with a positive log2FC in (a) (adj. p-value <1e-278). **c)** Same as

(b) but plotting the most differentially accessible region with a negative log2FC in (a) (adj. p-value<1e-278). **d)** Volcano plot for sequencer-associated differential signal for test case of 10 HiSeq4000 75 bp read samples against 10 NovaSeq6000 50 bp read samples. There are 146 DARs (p.adj<0.1), of which 75 have a large effect size (abs(log2FC)>1) and 71 have a small effect size (abs(log2FC)<1). **e)** Volcano plot for sequencer-associated differential signal from the same samples from (d), but the 75 bp HiSeq4000 reads trimmed down to 50 bp reads. There are 97 DARs (p.adj<0.1), of which 3 have a large effect size (abs(log2FC)>1) and 94 have a small effect size (abs(log2FC)<1). **f)** Plot of normalized chromatin reads as a function of trim length for region highlighted with a box in (d). **g)** Plot of normalized chromatin reads as a function of trim length for region highlighted with a circle in (d). **h)** Coverage and pileup plots of DAR from (e) when trimmed to 50 bp. Colored lines identify mismatches to reference genome: red – G, orange – C, dark blue – T, light blue – A. **i-j)** Comparison of differential accessibility signal between original 75bp/50bp analysis and trimmed 50bp/50bp analysis, with plots of log2FC (g) and -log10(padj) (h). Black line is a reference line with a slope of 1 and intercept of 0. Pearson correlations are given in bottom right corners. **k)** Boxplot comparing the transcription start site enrichment of samples that were sequenced on the HiSeq4000 and NovaSeq6000 (Student t-test p value<2e-16). **l)** Boxplot comparing the FRiP score of samples that were sequenced on the HiSeq4000 and NovaSeq6000 (Student t-test p value = 1e-11).

Supplementary Figure S4

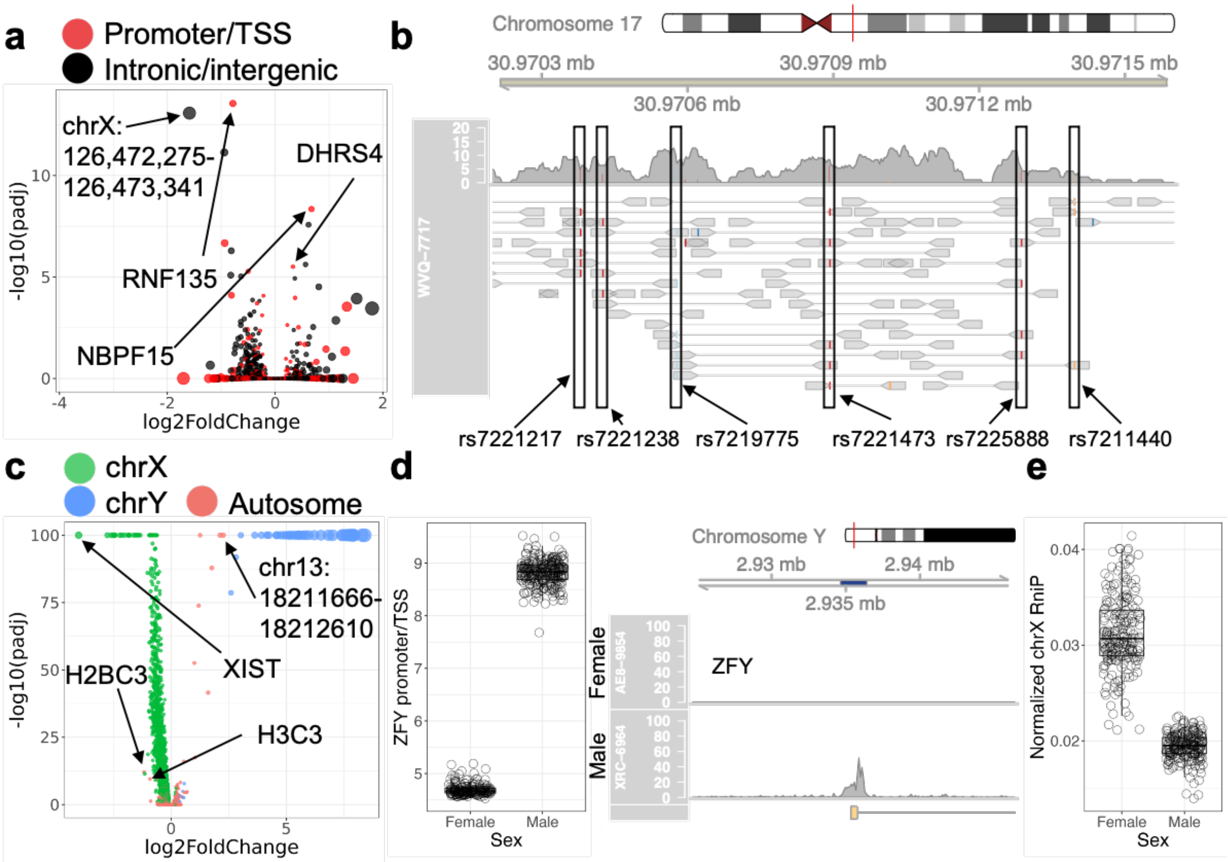


**Supplementary Figure S4. Clinical and demographic covariates extended data. a)** Volcano plot for ancestry-associated differential signal. **b)** Pileup and coverage plot of reads that mapped to the promoter/TSS of *RNF135*, a chromatin region whose accessibility was highly associated with ancestry for an individual of African ancestry. Boxes highlight consistent mismatches within mapped reads. **c)** Volcano plot for sex-associated differential signal. **d)** Normalized chromatin read counts (left) and example coverage plot (right) for *ZFY*, the most significant DAR on chromosome Y (adj. p-value < 1e-278). **e)** Reads **not** in peaks for chrX normalized to total reads across all chromosomes for female and male samples (p-value < 2e-16).

Supplementary Figure S5

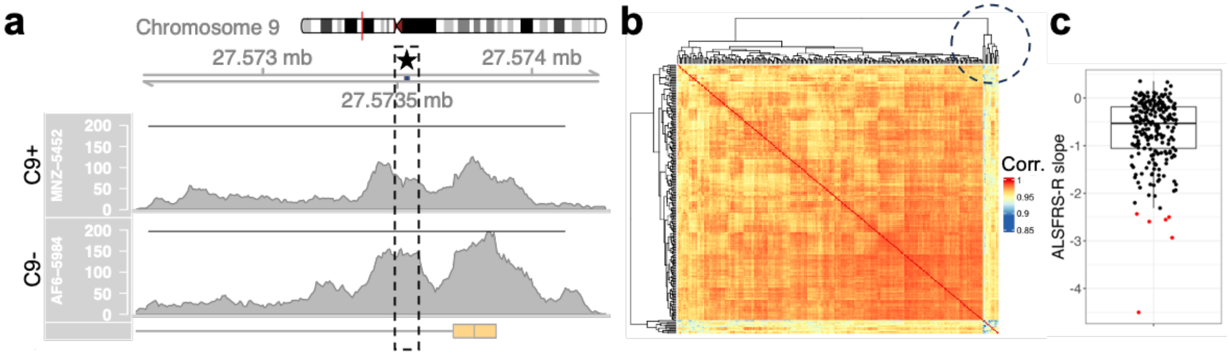


**Supplementary Figure S5. AALS ATAC-Seq ALS signals. a)** Example raw read coverage plot for the *C9orf72* TSS peak for a C9+ (repeat length of 274) and C9- ALS case. Region corresponding to the location of the repeat expansion is highlighted in dark blue in the genome axis scale under the star and surrounded by a box on the coverage plot. Black horizontal line is drawn at the same height in both coverage plots for reference. Note that differences in peak heights extend far beyond the region of the expansion. **b)** Correlation matrix of samples with ALSFRS-R slope using the normalized chromatin reads of regions that are within 2 kb of a TSS. Dendrogram was constructed using complete link hierarchical clustering with Euclidean distance between columns of the correlation matrix. Outlying samples are circled. **c)** Boxplot showing distribution of ALSFRS-R slopes after first round of filtering described in (a). Samples that were found to be outliers, as defined by being located more than 1.5 interquartile ranges from the first quartile, are colored in red.
